# Untargeted metabolomics reveals host responses and metabolites linked to host compatibility in Rafflesiaceae parasitism

**DOI:** 10.64898/2026.07.19.739411

**Authors:** Jeanmaire Molina, Rinat Abzalimov, Pride Yin, Adhityo Wicaksono, Faith Bernier, James Hill, Marco Bürger, Jun Wen, Susan K. Pell

## Abstract

Rafflesiaceae, including *Rafflesia, Sapria,* and *Rhizanthes*, are some of the rarest flowering plants in the world, known for producing the largest blooms on Earth. These holoparasites depend exclusively on *Tetrastigma* (Vitaceae) vines, yet the chemical basis for this host specificity is poorly understood, complicating conservation efforts. Untargeted negative-ion LC-MS metabolomics was used to profile Rafflesiaceae buds and seeds, infected *Tetrastigma* hosts associated with *Rafflesia lagascae, R. speciosa,* and *Sapria himalayana*, uninfected *Tetrastigma* hosts, and corresponding non-host species from the Philippines and Thailand. Python scripts used for data processing and visualization were developed with generative AI assistance and validated by the authors. Principal component analysis revealed distinct metabolomic profiles across samples, with significant shifts in host metabolites upon infection. Infected *Tetrastigma* tissues showed nominal enrichment in phenylpropanoid and gibberellin-related metabolites, suggestive of defense and hormonal pathway activation during *Rafflesia* infection, as well as citric acid, suggesting increased energy demand to support the parasite. Non-host *Tetrastigma* species showed nominal enrichment in metabolites putatively annotated as stilbenoids, resveratrol diglucosides, and condensed tannins such as procyanidin B2, compounds broadly associated with plant defense. *Rafflesia* seeds harbored fatty acids/oxylipins, while *Rafflesia* buds accumulated gallic acid derivatives, compounds also reported in galls. These findings support a chemically mediated model of host compatibility and parasitism, with implications for the ex-situ conservation of these endangered plants.

## 1. Introduction

Among the world’s flowering plants, *Rafflesia* stands out as one of the most extraordinary, hailed as “the greatest prodigy of the vegetable world” [1]. Famous for producing the largest flowers on Earth, some reaching up to a meter in diameter, *Rafflesia* species are obligate endoparasites found exclusively in Southeast Asia. *Rafflesia* is closely related to *Sapria* and *Rhizanthes*, together comprising the family Rafflesiaceae. All three genera are holoparasites that have lost their leaves, stems, and roots, and depend entirely on select species of *Tetrastigma* for water and nutrients [2, 3, 4]. These parasites spend most of their life embedded within host tissues and emerge only to flower, emitting an odor of rotting flesh to attract carrion flies in deceptive pollination.

Curiously, of the roughly 95 known species of *Tetrastigma*, which are widely distributed in subtropical and tropical regions of Asia and Australasia, only about 11 serve as hosts for *Rafflesia*, despite their overlapping geographic distributions [2]. These host species are scattered across the *Tetrastigma* phylogeny rather than forming a single clade. The factors governing *Rafflesia’s* highly selective host choice remain elusive. There is no evidence of strict co-speciation; individual *Tetrastigma* species can host multiple *Rafflesia* species that are not each other’s closest relatives [5]. Phylogenomic studies suggest this parasitic relationship is ancient, dating back to the Cretaceous period, well before *Tetrastigma* itself is hypothesized to have emerged (∼40 Mya), based on evidence of horizontal gene transfer (HGT) from parasite to host [6, 7]. This suggests that ancestral *Rafflesia* may have parasitized other Vitaceae members such as *Ampelopsis* [7]. This long evolutionary history has likely enabled *Rafflesia* (and its close relatives *Rhizanthes* and *Sapria*) to discard the chloroplast genome entirely, which reflects their total reliance on the host with complete loss of photosynthetic capacity [7, 8].

Unfortunately, most *Rafflesia* species today are endangered due to habitat loss from deforestation [9]. Their already precarious status is compounded by biological and ecological mysteries: the factors governing host specificity remain unknown; *Rafflesia* exhibits extremely high bud mortality (>90%); and its seed ecology and germination remain enigmatic [11]. Ex-Situ conservation efforts have been largely unsuccessful. Numerous in vitro and in vivo germination trials using various plant growth regulators have failed to induce *Rafflesia* seed germination [11, 12, 13]. Grafting *Rafflesia*-infected *Tetrastigma* onto uninfected hosts yielded promising results, producing multiple blooms between 2004 and 2010 at Bogor Botanic Garden (BBG) [11]. More recently, *Rafflesia* buds emerged from grafted, infected *Tetrastigma* cuttings collected from the Philippines and cultivated at the U.S. Botanic Garden, marking the first successful propagation of *Rafflesia* in the Western Hemisphere and a major breakthrough in ex situ conservation [14]. A full understanding of *Rafflesia’s* enigmatic life cycle, including the biology and chemical ecology of its dispersal and host infection, would better enable cultivation for ornamental, educational, and conservation purposes.

Microbial profiling has revealed that *Rafflesia speciosa* seeds harbor endophytes similar to those in their *Tetrastigma* hosts, suggesting possible microbiota transmission during infection [15]. In other parasitic plants, such as *Striga*, altered soil microbiomes have been shown to disrupt parasitism by impairing haustorium formation [16]. Similar microbiome effects have been observed in the plant parasites *Langsdorffia* and *Orobanche*, where host-associated microbes influence parasite success [17, 18]. These findings point to a likely role of microbial interactions in shaping *Rafflesia*-*Tetrastigma* compatibility.

Chemical ecology also offers key insights. Transcriptomic profiling of *Rafflesia* seeds has identified genes linked to plant growth regulators that may facilitate germination [19]. LC-MS-based metabolomics revealed the presence of benzylisoquinoline alkaloids, defense-related compounds in *Tetrastigma* shoots, with reduced levels in infected tissues, indicating host chemical changes during parasitism [20]. Such results underscore the potential of metabolomics to illuminate host selectivity and infection dynamics. In a recent study, Molina et al. [21] combined metagenomics and metabolomics to explore functional links between microbes and metabolites in this system. They found associations between bacterial families (e.g., Microbacteriaceae, Nocardioidaceae) and elevated polyphenols, especially gallic acid derivatives, which may create a favorable chemical environment for *Rafflesia*. Bacteria capable of degrading complex carbon compounds were enriched in *Rafflesia* buds, while an unidentified Saccharimonadales group was enriched in the host. The fatty acid amide docosenamide was detected in both buds and host tissues, suggesting a role in facilitating parasitic infection. Meanwhile, coumarins, which are abundant in non-host *Tetrastigma* species, may act as allelopathic defenses against parasitism. The accumulation of gallic acid derivatives, adenine, and gall-associated bacteria in *Rafflesia* buds seems to support the hypothesis that *Rafflesia* acts like plant galls, manipulating host tissue to favor parasite development [21].

Building on these findings, the present study applies computational tools to process, visualize, and summarize LC-MS data. Generative AI was used in a limited capacity to assist with drafting and refining Python code, with all scripts and outputs reviewed and validated by the authors. Recent advances in artificial intelligence and large language models (LLMs) such as ChatGPT have expanded their applications in analytical chemistry and metabolomics by facilitating statistical coding assistance, multivariate visualization, literature synthesis, and exploratory data interpretation [22, 23, 24]. In general, AI-assisted workflows are increasingly used as supplementary computational aids, although their outputs require careful validation, domain expertise, and author oversight to ensure analytical accuracy and reproducibility [25].

While the previous study [21] employed LC-MS in positive ion mode, the present study analyzes the same samples in negative mode (ESI-), which is optimal for detecting acidic molecules (e.g., phenols, carboxylic acids, certain flavonoids, and polyphenols) and complements the prior results that favored basic nitrogen-containing compounds (e.g., alkaloids, docosenamide) [21]. The present study also included samples of *Ampelopsis*, which was hypothesized to be an ancestral Rafflesiaceae host based on phylogenomic evidence of horizontally transferred genes as DNA fossils [7]. If *Ampelopsis,* the first diverged extant lineage in Vitaceae [26, 27], was an ancient host, we predict that this genus will share metabolomic features with *Tetrastigma* hosts. We also included samples of *Rafflesia speciosa* seeds and juvenile stems/leaves of *T. harmandii,* the host of *R. speciosa*, to determine organ-specific changes in *Rafflesia* development and host chemistry, respectively, that may shed light on why *R. speciosa* only infects and emerges in host roots and not stems/leaves of the same host.

The primary goal of the present study is to better understand the chemical ecology of host and non-host *Tetrastigma* species and to profile metabolomic changes across the life cycle of *Rafflesia*, from seeds to buds, particularly in *R. speciosa*. To facilitate the analysis of these high-dimensional LC-MS datasets, we leveraged AI-assisted tools to support data processing, statistical scripting, visualization, and exploratory interpretation, while metabolite annotations were generated using curated spectral and natural-product databases and reviewed by the authors. Through this integrative approach, we aim to identify key metabolites associated with host compatibility, parasitism, and *Rafflesia* development, thereby providing insights that may inform more effective strategies for the ex situ conservation of this botanical marvel.

## 2. Materials and Methods

### 2.1 Plant material preparations

We sampled Rafflesiaceae buds and seeds, infected *Tetrastigma* hosts associated with *Rafflesia lagascae, R. speciosa,* and *Sapria himalayana,* uninfected *Tetrastigma* hosts, and corresponding non-host species from Iloilo and Camarines Norte, Philippines, and Queen Sirikit Botanic Garden, Mae Rim, Chiang Mai, Thailand. Most materials were identical to those used in Molina et al. [21], with additional samples in the present study, including *R. speciosa* seeds (n = 2), uninfected juvenile stems/leaves of *T. harmandii* (n = 2), and four *Ampelopsis* species: *A. bodinieri*, *A. glandulosa*, *A. delavayana*, and *A. cordata*, collected from the Smithsonian Gardens greenhouses.

Two *Rafflesia-Tetrastigma* systems were sampled in the Philippines: *R. speciosa* from Iloilo (ILO, buds n = 8) and *R. lagascae* from Camarines Norte (CAM, buds n = 2). *Rafflesia speciosa* parasitizes *Tetrastigma harmandii* and *T. cf. magnum*, whereas *R. lagascae* infects *T. loheri* and *T. sp. A*. *Tetrastigma* individuals bearing visible *Rafflesia* buds were classified as infected (ILO = 9; CAM = 2). Plants visibly lacking buds were classified as uninfected hosts (ILO = 10; CAM = 2), though dormant endophytic infections cannot be ruled out. Other *Tetrastigma* species, *T. scariosum*, *T. papillosum*, and a reddish morph of *T. aff. loheri,* have never been observed to host *Rafflesia* and are considered non-hosts (ILO n = 4; CAM n = 2). Additional samples of *Sapria himalayana* buds (n = 2) and their *Tetrastigma* hosts (n = 2 infected: *T. obovatum*, *T. cruciatum*; n = 2 uninfected: *T. cauliflorum*) were collected from Queen Sirikit Botanic Garden, Thailand (THAI).

Fifty-seven plant tissue samples were analyzed: *R. speciosa* seeds (n = 2), *Rafflesia/Sapria* buds (n = 12), infected *Tetrastigma* cuttings (roots and stems) within 5 cm of buds (n = 13), uninfected host samples (including juvenile stems and leaves, n = 16), root pieces of non-host *Tetrastigma* (n = 6), and *Ampelopsis* spp. (n = 8). Each sample was analyzed in duplicate, for a total of 114 samples. All materials were submitted for metabolite profiling at the Advanced Science Research Center, City University of New York (CUNY). Samples were standardized to a concentration of 0.05 mg μL ¹ during methanol extraction. Extracts were reconstituted in 0.3 mL of 1:1 (v/v) methanol/water prior to LC-MS injection.

### 2.1 LC-MS analysis

LC-MS profiling was performed on a maXis-II UHR-ESI-QqTOF (Bruker Daltonics, USA) mass spectrometer coupled to an UltiMate 3000 UHPLC (Thermo Scientific, USA) system using a reverse-phase C18 column. The LC gradient was set as follows: 2% solvent B (acetonitrile with 0.15% formic acid) and 98% solvent A (water with 0.15% formic acid) for the first 2 minutes, followed by an increase to 40% solvent B over 20 minutes, to 98% solvent B over the next 10 minutes, and held at 98% solvent B for an additional 10 minutes. Mass spectrometry data were acquired over an m/z range of 50–1300 in negative-ion electrospray ionization mode.

### 2.3 Data processing

Raw data were processed using MetaboScape 2025 software (Bruker Inc, USA). Features were filtered to remove blanks/background and normalized to the internal standard. Compound annotation was performed using Bruker MetaboBASE Personal Library 3.0, LipidBlast2022 at the MassBank of North America (MoNA) [28] (https://mona.fiehnlab.ucdavis.edu), and the COlleCtion of Open NatUral producTs or COCONUT [29, 30] (https://coconut.naturalproducts.net/). Only likely natural products (phenolics, flavonoids, glycosides, terpenoids, tannins) were retained; synthetic motifs were excluded.

The complete metabolomics matrix and associated sample metadata are provided as Supplementary Dataset S1 and S2, respectively.

### 2.4 Data analysis

Metabolomic data were analyzed to evaluate chemical differentiation among Rafflesiaceae tissues, infected and uninfected *Tetrastigma* hosts, non-host *Tetrastigma* species, aerial host tissues, *Rafflesia speciosa* seeds, and *Ampelopsis* species. Normalized LC-MS feature intensities were used for all downstream analyses. Prior to visualization and statistical testing, duplicate metabolite annotations were consolidated by averaging intensities for rows with the same metabolite name or feature label where appropriate. Negative values were set to zero for abundance-based composition plots. All statistical analyses and visualizations were performed in Python. Portions of the coding workflow were assisted by ChatGPT GPT 5.2 and later 5.5 (OpenAI, San Francisco, CA, USA), using a high reasoning-effort (“extended thinking”) setting, which was used to generate and refine Python scripts for data handling, statistical calculations, and figure generation. All AI-assisted scripts, outputs, and visualizations were reviewed, tested, and executed by the authors to ensure accuracy, reproducibility, and compliance with journal guidelines on the responsible use of generative artificial intelligence. The generated scripts are stored in GitHub (see Data Availability Statements). In addition, the manuscript text, figures, and underlying statistical analyses were subsequently reviewed and revised with the assistance of Claude Sonnet 5 (Anthropic, San Francisco, CA, USA), which was used to check the accuracy and reproducibility of the Python scripts described above. All Claude-assisted edits were reviewed and approved by the authors prior to submission.

Principal component analysis (PCA) was used to assess broad metabolomic separation among sample groups. PCA was performed on normalized metabolite abundance matrices using Python, and samples were visualized in three-dimensional ordination space using the first three principal components. Separate PCA analyses were conducted for: 1) all samples, including Rafflesiaceae tissues, *Tetrastigma*, and *Ampelopsis*; 2) host, non-host, and *Ampelopsis* samples (parasitic tissues excluded); and 3) samples pooled by biological interaction status: infected hosts, uninfected hosts, non-hosts, seeds, buds, and uninfected aerial stem/leaf tissue, combined across localities to emphasize infection status, host compatibility, and developmental stage rather than site-level differences. Sample groups were distinguished by color and marker shape, and confidence ellipsoids were included for groups with sufficient replication.

Next, the differential metabolite enrichment was assessed using log_2_ fold-change (log_2_FC) analyses calculated from normalized mean intensities between biologically relevant contrasts. Comparisons included infected versus uninfected *Tetrastigma* hosts pooled across regions, as well as locality-specific comparisons between non-host and infected *Tetrastigma* tissues from Camarines Norte and Iloilo. Statistical significance was evaluated using two-sided Welch’s *t*-tests. Metabolites with *p* < 0.05 and log_2_FC > 1 were considered enriched. Significance scatterplots and log_2_FC bar plots were used to visualize infection-associated and non-host-enriched metabolites, with selected natural-product annotations labeled.

To examine metabolomic shifts across the *Rafflesia speciosa* life cycle and associated host tissues, group-level mean abundances were calculated for seeds, buds, infected host tissues, uninfected host tissues, aerial stem/leaf tissues, and non-host roots. For stacked bar plots, feature intensities were normalized by total-sum scaling within each group to show the relative contribution of the most abundant compounds and major chemical classes.

Heatmaps were used to visualize relative enrichment of selected natural metabolites across infected host roots, uninfected host roots, uninfected aerial stem/leaf tissues, and non-host roots in the *R. speciosa*-*Tetrastigma* system. Metabolites were selected based on annotation confidence, natural-product status, and ecological relevance to host defense, infection, primary metabolism, or parasitic development. Mean metabolite abundances were standardized using row-wise Z-score normalization, allowing relative enrichment or depletion of each metabolite to be compared across tissue types. Heatmaps were generated in Python using Matplotlib with a diverging color scale, where positive Z-scores indicate relative enrichment and negative Z-scores indicate relative depletion.

## 3. Results

### 3.1 Distinct metabolomic profiles across parasitic, host, and non-host tissues

Untargeted LC-MS metabolomics in negative ion mode revealed broad chemical differentiation among Rafflesiaceae tissues, infected and uninfected *Tetrastigma* hosts, non-host *Tetrastigma* species, and *Ampelopsis.* Principal component analysis (PCA) of all samples showed separation associated with each sample’s biological role in the host-parasite system (parasitic tissue, infected host, uninfected host, non-host, or *Ampelopsis*) as well as tissue/source group, with the first three components explaining 26.3% of cumulative variance (PC1: 9.6%, PC2: 9.0%, PC3: 7.7%). Rafflesiaceae buds and *R. speciosa* seeds separated from most host and non-host vine tissues, while *Ampelopsis* occupied a distinct region of multivariate space. *Tetrastigma* hosts (infected and uninfected) and non-hosts showed partial separate clustering, although some overlap was observed among vine tissues (Fig. 1).

**Fig. 1.**
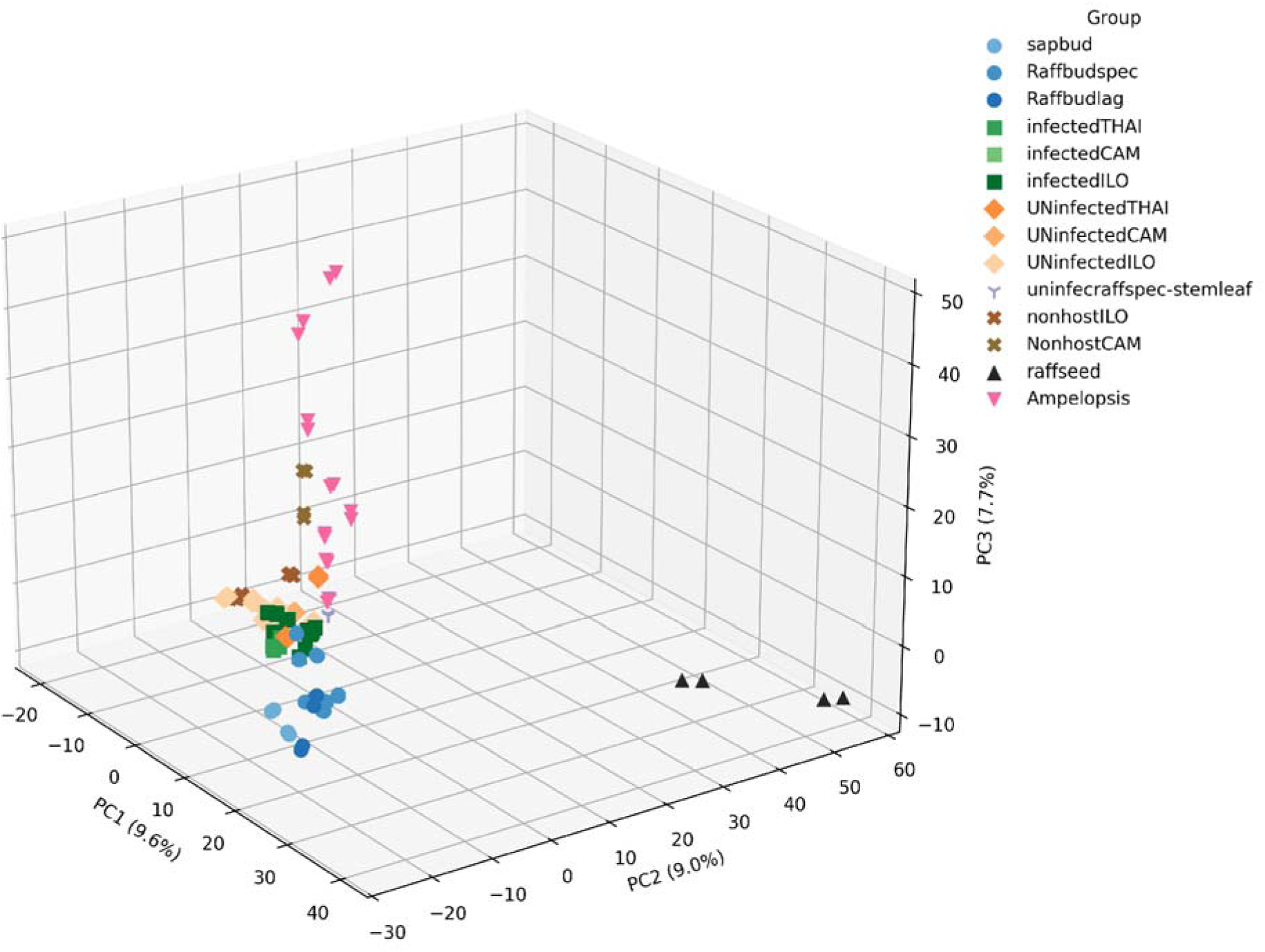
Principal component analysis (PCA) of metabolite profiles across *Rafflesia*, *Tetrastigma*, and *Ampelopsis* sample groups. The 3D PCA plot displays the first three principal components, which together explain 26.3% of the cumulative variance (PC1: 9.6%, PC2: 9.0%, PC3: 7.7%). Each point represents an LC-MS technical run, with two runs per biological sample, colored by sample group and shaped by sample category, reflecting tissue type and host-parasite relationship: circles for *Rafflesiaceae* buds, squares for infected hosts, diamonds for uninfected hosts, Xs for non-hosts, triangles for *Rafflesia* seeds (black), and inverted triangles for *Ampelopsis*. The PCA shows broad chemical differentiation among parasitic tissues, host tissues, non-host species, and *Ampelopsis*, with partial overlap among several vine tissue groups.

When parasitic tissues (i.e., *Rafflesia speciosa* seeds and *Rafflesia/Sapria* buds) were excluded, PCA of host and non-host vine tissues showed partial clustering by infection status and tissue type. Infected *Tetrastigma* samples from the Philippines and Thailand occupied a broadly similar region of PCA space despite geographic separation, although they partially overlapped with uninfected host and non-host samples. Juvenile stems and leaves of *T. harmandii* were chemically distinct from infected root tissues of the same host species, separating primarily along PC2 and PC3. Non-host *Tetrastigma* samples tended to cluster separately from infected hosts within the same locality, while *Ampelopsis* formed a comparatively distinct group. Ellipsoids indicate group-level dispersion where sufficient samples were available (Fig. 2).

**Fig. 2.**
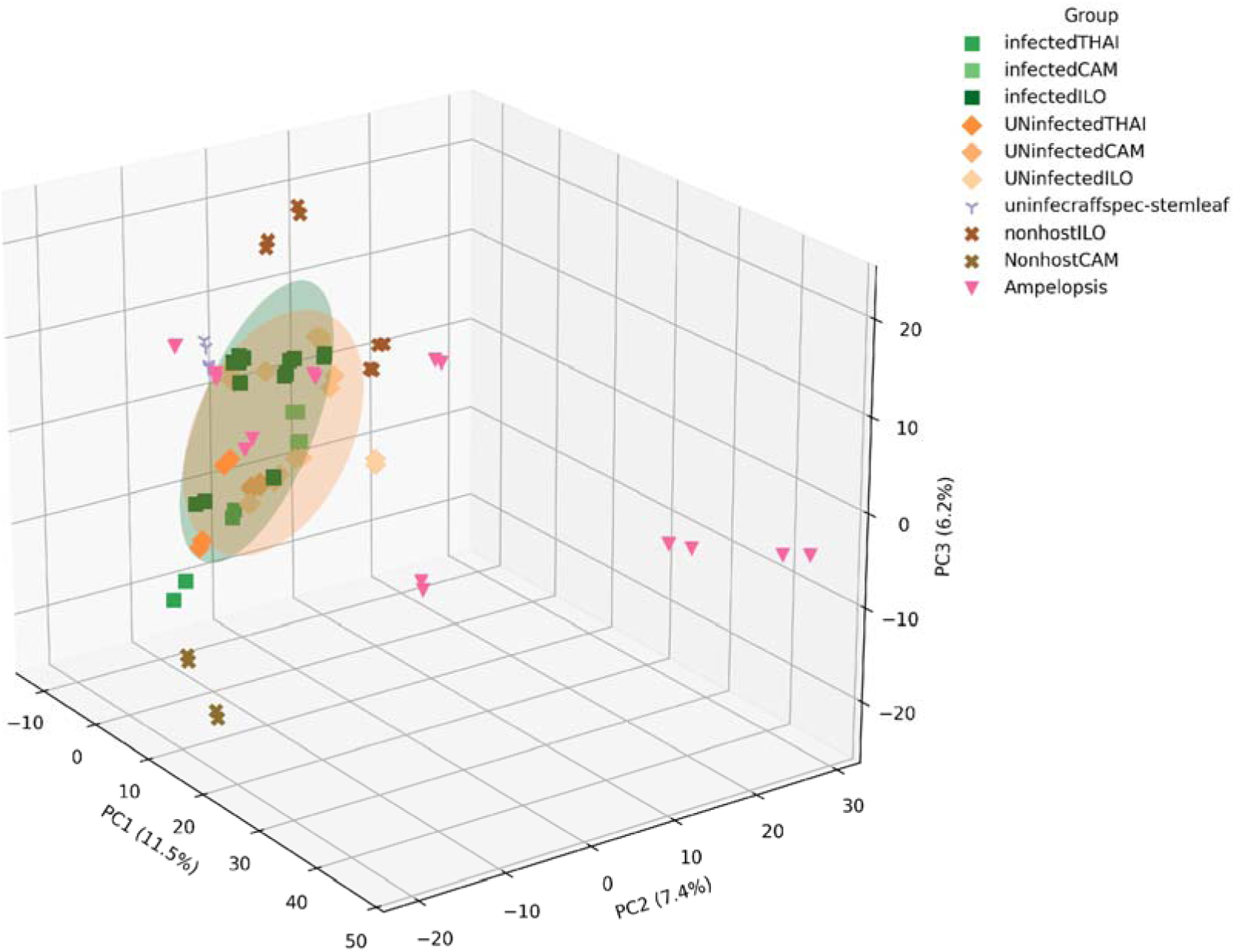
Principal component analysis (PCA) of metabolite profiles across infected, uninfected, and non-host *Tetrastigma* species, as well as *Ampelopsis* with Rafflesiaceae sample groups removed. 3D PCA plot shows the first three principal components, which together explain 25.1% of cumulative variance (PC1: 11.5%, PC2: 7.4%, PC3: 6.2%). Each point represents one LC-MS run, with two technical runs per biological sample. Groups are distinguished by color and marker shape. Ellipsoids represent approximately 68% covariance regions for groups with sufficient replication (based on Mahalanobis distance in PC1-PC3 space) and were pooled across localities rather than fit separately per site. Infected hosts (squares, *Tetrastigma* pooled across Thailand, Camarines Norte, and Iloilo) are enclosed within a single green ellipsoid, while uninfected hosts (diamonds, pooled across the same three localities) are enclosed within a single orange ellipsoid; non-host *Tetrastigma* (cross marks) are shown without an ellipsoid given smaller per-locality sample sizes. The two ellipsoids overlap partially, consistent with chemical divergence between infected and uninfected hosts. Uninfected juvenile shoots and leaves of *Rafflesia speciosa*’s host (lilac tripod/Y-markers) are visually separated from the infected host roots (infectedILO, dark green squares). Non-host *Tetrastigma* species (nonhostILO and NonhostCAM, cross marks) are visually separated from infected species (infectedILO and infectedCAM) within the same locality (CAM, ILO). *Ampelopsis* is represented with pink triangle markers. This visualization illustrates chemical differences among host-compatibility groups.

Grouping samples by biological interaction status (e.g., infection status, host compatibility, and developmental stage, pooled across localities) further emphasized broad metabolomic separation among Rafflesiaceae buds, infected hosts, uninfected hosts, non-hosts, seeds, and uninfected stem/leaf tissues. Rafflesiaceae buds and seeds occupied distinct regions of PCA space, while infected, uninfected, and non-host vine tissues showed partial clustering with some overlap. Infected hosts and non-host *Tetrastigma* samples were generally separated, although a few samples occupied nearby regions of multivariate space (Fig. 3).

**Fig. 3.**
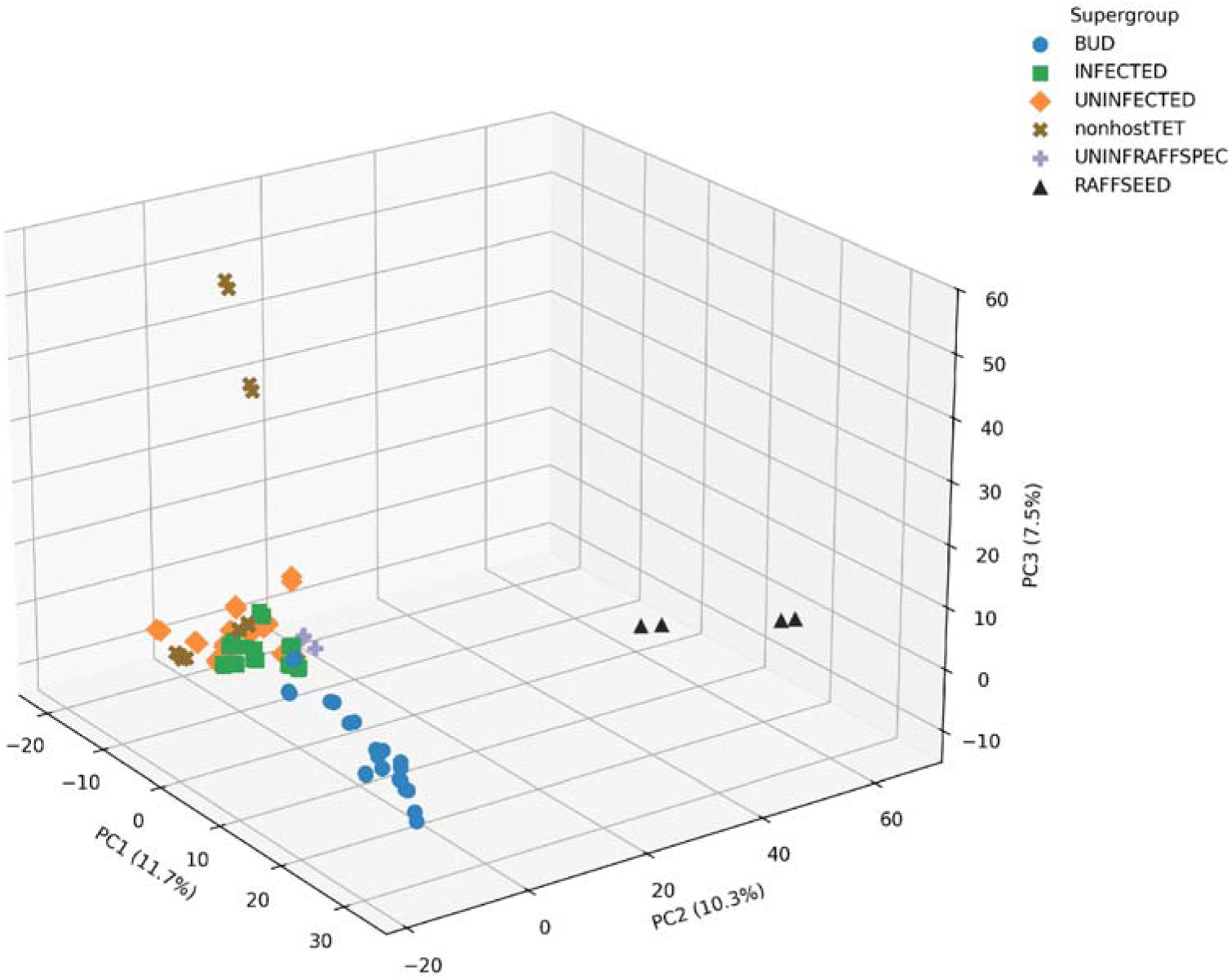
Principal component analysis (PCA) of metabolite profiles from *Tetrastigma* and parasitic associations (e.g. *Rafflesia speciosa* seed, *Rafflesia* and *Sapria* buds). Each point represents one LC-MS run, with two technical runs per biological sample, projected along the first three principal components, which together explain 29.5% of the cumulative variance (PC1: 11.7%, PC2: 10.3%, PC3: 7.5%). Sample groups are color-coded and symbolized as follows: BUD (blue circles: sapbud, Raffbudspec, Raffbudlag), INFECTED (green squares: infectedTHAI, infectedCAM, infectedILO), UNINFECTED (orange diamonds: UNinfectedTHAI, UNinfectedCAM, UNinfectedILO), non-hostTET (brown Xs: nonhostILO, NonhostCAM), RAFFSEED (black triangles), and UNINFRAFFSPEC (purple triskele-style markers). The exclusion of *Ampelopsis* enhances the focus on biologically relevant *Tetrastigma* lineages, revealing chemical clustering based on infection status and parasitic interaction.

### 3.2 Infection-associated metabolites conserved across regions

The significance scatterplot (Fig. 4) identified five metabolites nominally enriched in infected *Tetrastigma* tissues (uncorrected *p* < 0.05, log_2_FC > 1): Nb-trans-feruloylserotonin glucoside, gibberellin A93, ellagic acid, 4-methoxycinnamic acid, and quercetin-3-O-deoxyhexosyl(1→2)pentoside. None of these five metabolites remained significant after Benjamini–Hochberg FDR correction (*q* < 0.05), so they are best treated as candidate markers warranting targeted validation rather than confirmed infection-associated compounds. These compounds nonetheless represent multiple chemical classes, including phenolic acid/phenylpropanoids, flavonoids, glycosides, and phytohormones.

**Fig. 4.**
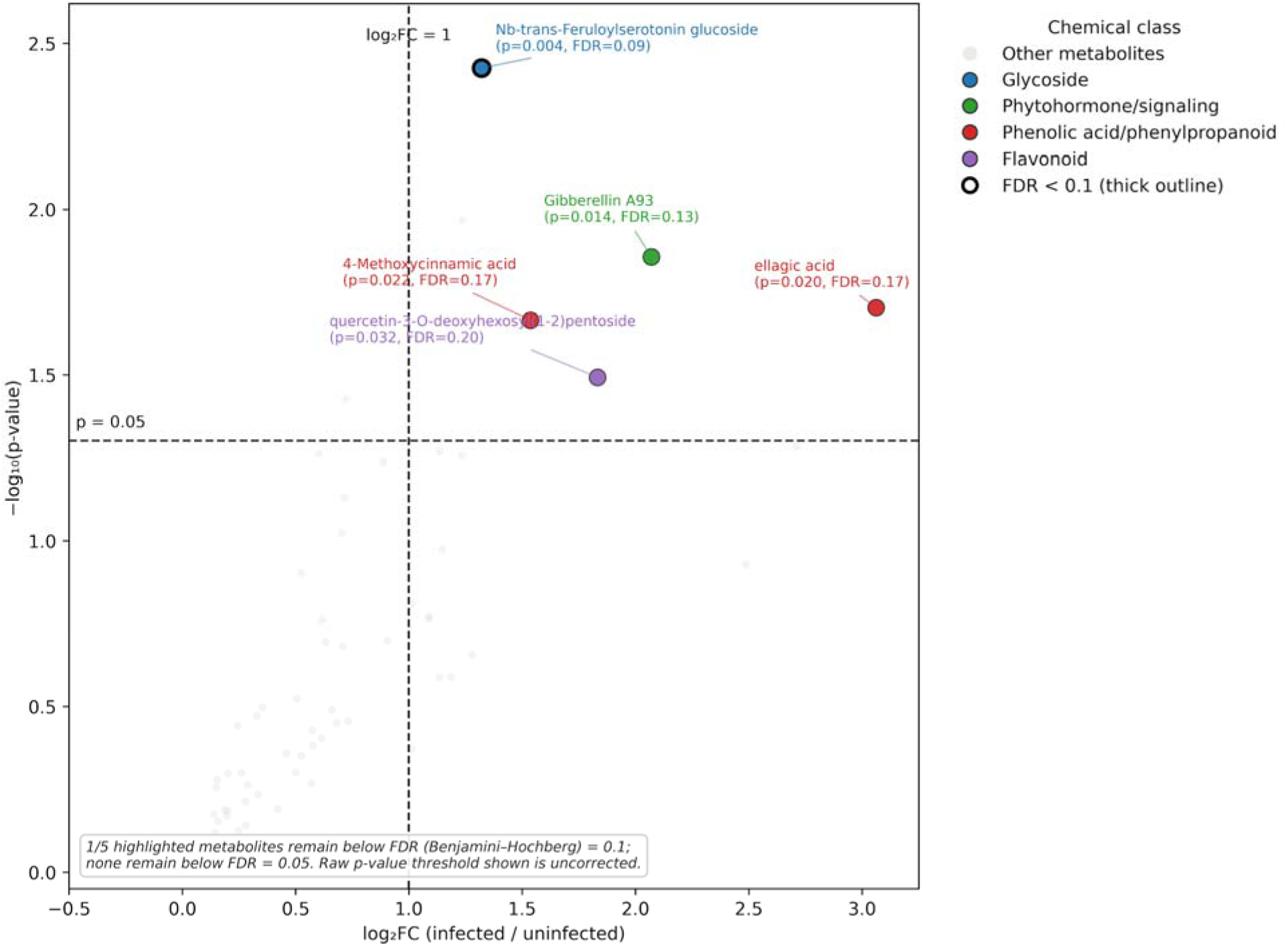
Differential abundance of natural metabolites in infected versus uninfected Tetrastigma hosts. Significance scatterplot showing metabolite enrichment in infected *Tetrastigma* tissues relative to uninfected tissues pooled across Thailand/THAI, Camarines Norte/CAM, and Iloilo/ILO. The x-axis represents log_2_FC (infected/uninfected), and the y-axis shows −log_10_(p-value) from two-sided Welch’s t-tests. Metabolites were filtered to retain natural, non-questionable annotations prior to analysis. Labels report both the raw p-value and the Benjamini– Hochberg FDR-adjusted q-value for each highlighted metabolite; a heavier point outline indicates FDR < 0.10. Five highlighted natural-product annotations showed nominal infection-associated enrichment under the uncorrected p-value threshold, but none remained significant after Benjamini–Hochberg correction at FDR < 0.05. Dashed lines indicate the uncorrected fold-change threshold (log_2_FC = 1) and significance threshold (*p* = 0.05). To improve visualization, metabolites with log_2_FC < −0.5 were omitted from the plot. None of the labeled metabolites remained significant after FDR correction (*q* < 0.05); these should therefore be interpreted as nominal, exploratory candidate markers rather than statistically confirmed markers of Rafflesiaceae-infected host tissues.

### 3.3 Non-host *Tetrastigma* species exhibit distinct chemical defenses

Non-host *Tetrastigma* species displayed metabolomic profiles distinct from infected hosts (Fig. 5). Two compounds remained significant after Benjamini–Hochberg FDR correction in at least one locality: Caffeoylmalic acid and Epicatechin-(4β→8)-gallocatechin, the latter via distinct ion features significant at CAM and ILO, respectively, rather than the same feature replicating across both sites. Other candidates showing nominal (uncorrected p < 0.05) but not FDR-corrected enrichment included stilbenoids (e.g., (Z)-resveratrol 3,4′-diglucoside) and additional flavonoid/tannin derivatives ((−)-Epigallocatechin 3-p-coumaroate; 2′,7-dihydroxy-4′-methoxy-8-prenylflavan diglucoside). Given the small sample size at CAM in particular (n = 2 per group), these should be treated as candidate rather than confirmed markers of non-host chemical defense.

**Fig. 5.**
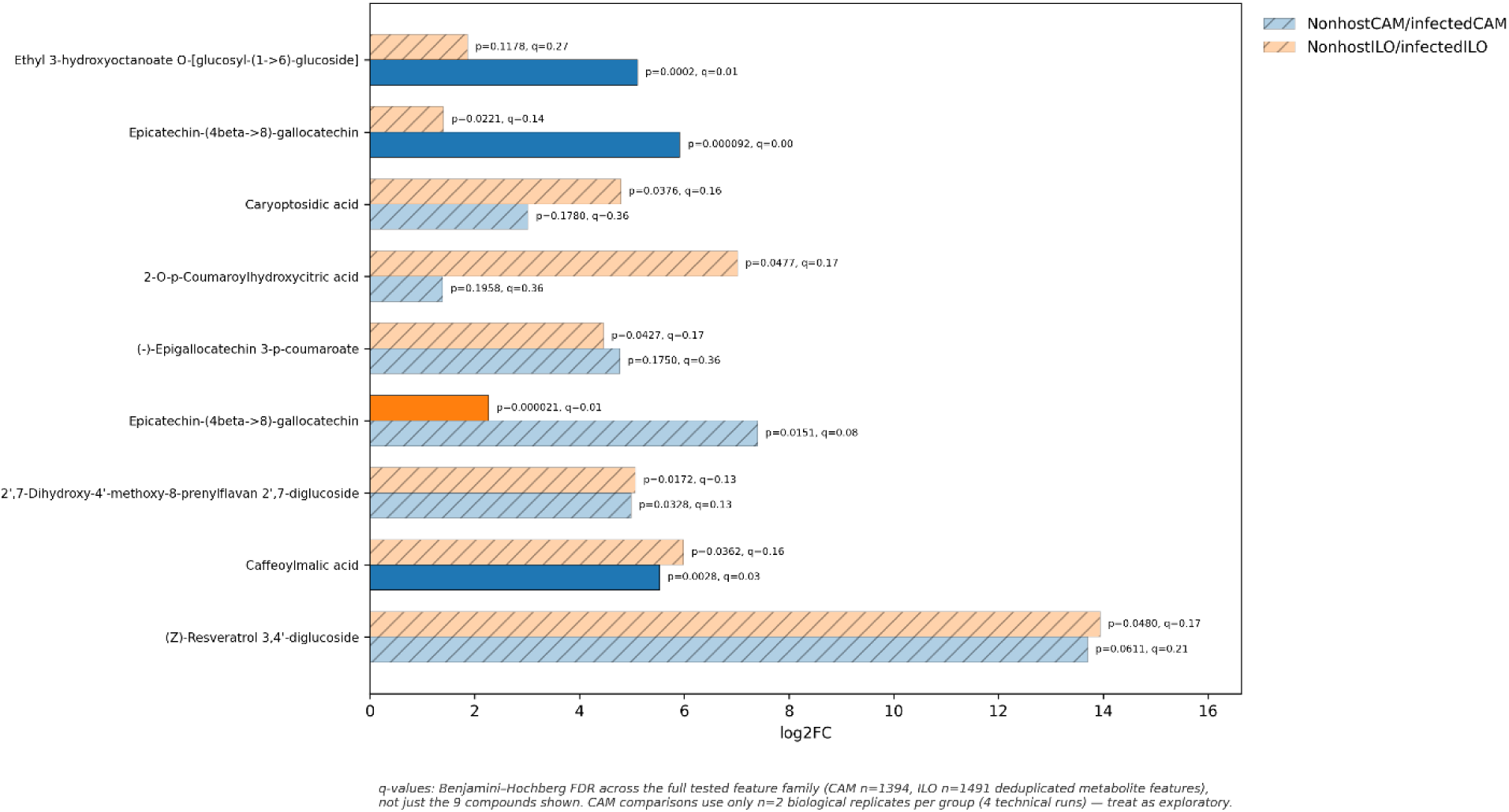
Log_2_FC change in compound abundance between non-host and infected *Tetrastigma* tissues from two geographic locations (CAM and ILO only, since non-host *Tetrastigma* spp. were not found and collected in THAI). Bar plots display log_2_FC values, with both the raw Welch’s t-test p-value and the Benjamini–Hochberg FDR-adjusted q-value annotated for each comparison. Because only 9 compounds are shown out of the full tested feature set, q-value were calculated across the complete family of metabolite features tested per locality (CAM n = 1,394; ILO n = 1,491 deduplicated features), not just the 9 shown, to avoid inflating apparent significance through post-hoc selection. Bars with FDR ≥ 0.05 are hatched. Metabolites were selected for display based on log_2_FC magnitude and/or nominal statistical support in at least one locality; most did not remain significant after correction, and the Camarines Norte (CAM) comparisons in particular rest on only two biological replicates per group and should be interpreted as exploratory. Several feature-level comparisons retained FDR support after correction, although enrichment patterns varied by locality and should be interpreted cautiously because some comparisons rely on limited biological replication.

### 3.4 Metabolomic shifts across *Rafflesia speciosa* life stages and associated host tissues

Metabolite composition differed markedly across *Rafflesia speciosa* seeds, buds, host tissues, and non-host species. Seeds were dominated by fatty acids and oxylipins, whereas buds accumulated polyphenolic compounds, particularly gallic acid derivatives and epicatechin-based tannins. Infected *Tetrastigma* tissues showed elevated citric acid relative to uninfected hosts and non-hosts, while non-host species were enriched in condensed tannins such as procyanidin B2 (Fig. 6).

**Fig. 6.**
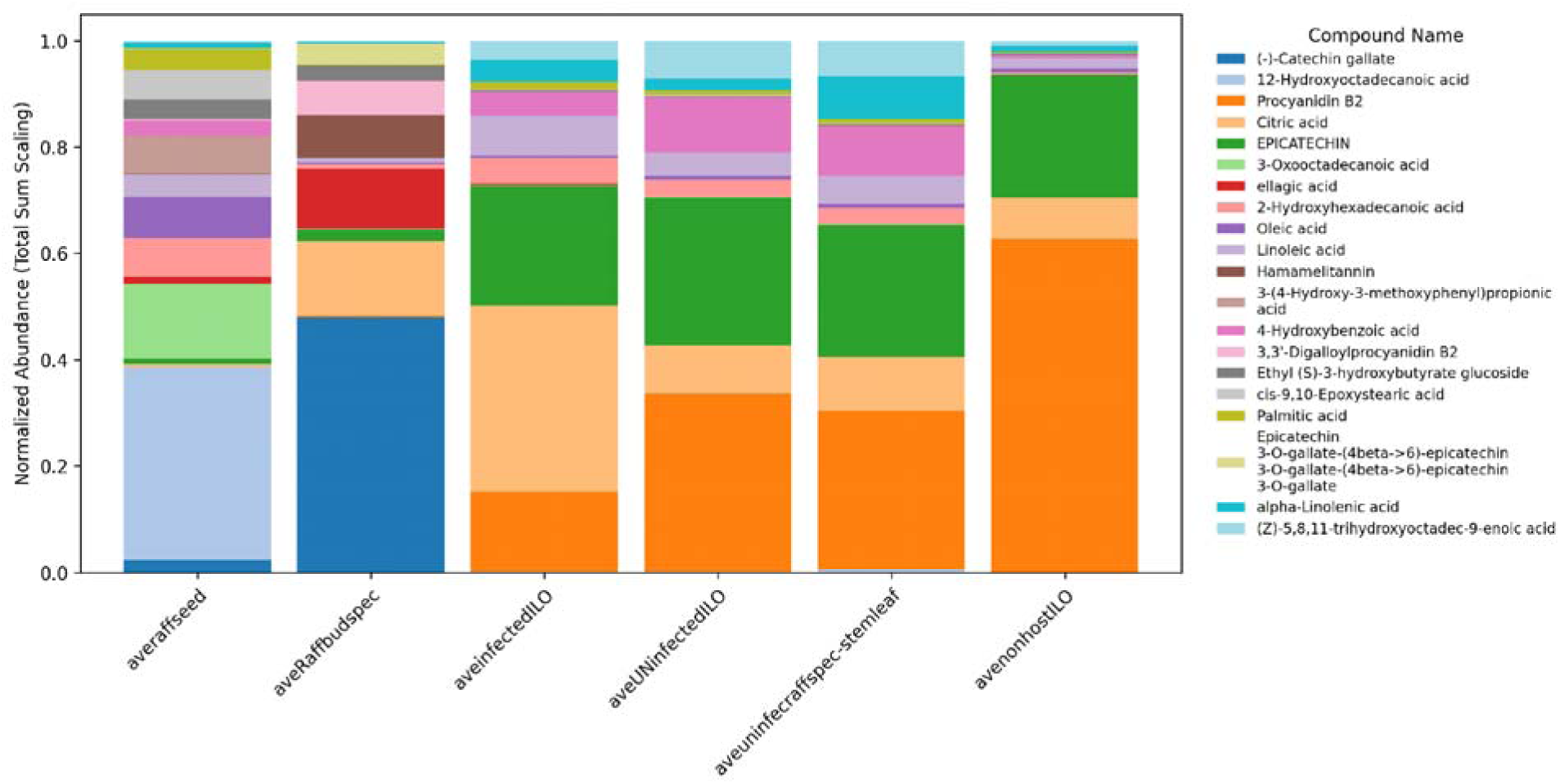
Relative metabolite composition across *Rafflesia speciosa* seeds (RAFFSEED), buds (RAFFBUD), infected and uninfected *Tetrastigma* host tissues, and non-host species from the same Philippine locality (ILO). Stacked bar plot of the top 20 most abundant compounds (normalized by total sum scaling, negative values set to zero), highlighting tissue-specific enrichment such as fatty acids/oxylipins in *Rafflesia* seeds, tannins (e.g., epicatechin derivatives) in buds, elevated citric acid in infected *Tetrastigma*, and procyanidin B2 in non-host species. These profiles underscore developmental stage-specific metabolism in *Rafflesia* and chemical convergence with host tissues during infection.

The heatmap (Fig. 7) revealed distinct tissue-specific metabolic signatures across *Tetrastigma* samples. Infected host roots were characterized by elevated levels of organic acids (citric acid, L-malic acid) and flavonoids (eriodictyol-7-O-glucoside and (-)-epicatechin-3′-O-glucuronide). Uninfected roots were depleted in these compounds but contained slightly more of the long-chain fatty acid, 2-hydroxyhexadecanoic acid. Aerial stem and leaf tissues of host species were enriched in fatty acids (α-Linolenic acid), flavonoids (apigenin 7-O-glucoside, isovitexin, etc.), and phenylpropanoids (trans-o-Coumaric acid 2-glucoside, 5-O-Feruloylquinic acid). In contrast, non-host roots accumulated defense-associated metabolites, including proanthocyanidins (procyanidins, cinnamtannins), phenylpropanoids (ferulic acid, caffeic acid conjugates, 5-hydroxy-6-methoxycoumarin 7-glucoside, etc), terpenoids (oleanolic acid, isolimonic acid), and were generally deficient in most of the metabolites described as enriched in host species.

**Fig. 7.**
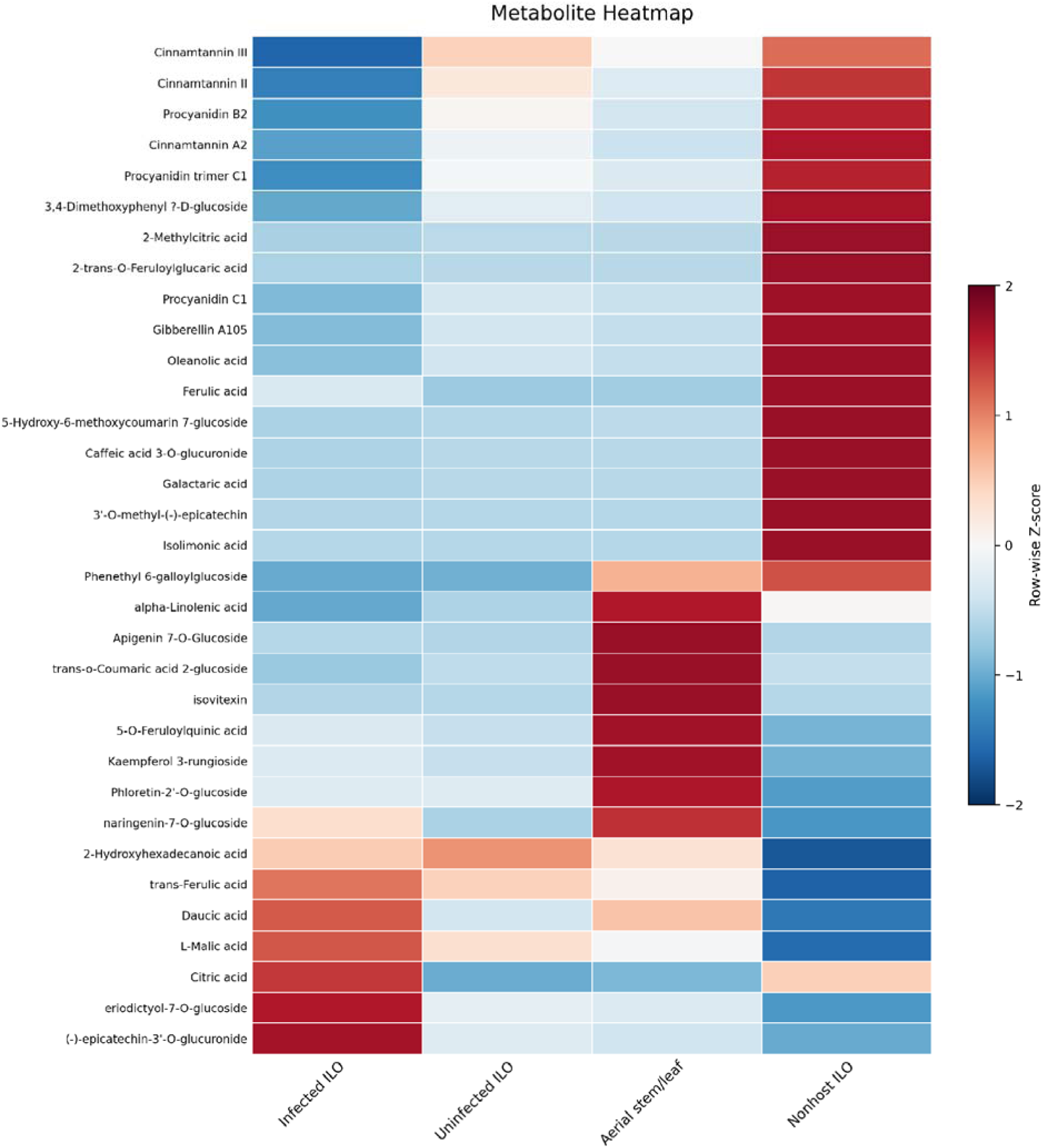
Heatmap showing the relative abundance of selected natural metabolites across infected host roots (Infected ILO), uninfected host roots (Uninfected ILO), uninfected aerial stem/leaf tissues, and non-host roots (Non-host ILO) in the *Rafflesia speciosa-Tetrastigma* system. Metabolites were selected from the untargeted LC-MS dataset based on their ecological relevance and confidence of annotation, and values were standardized using row-wise Z-score normalization to facilitate comparisons among tissue types. Positive Z-scores (orange-red) indicate relative enrichment, whereas negative Z-scores (blue) indicate relative depletion relative to the mean abundance of each metabolite across tissues.

## 4. Discussion

This study provides novel insights into the chemical ecology of *Rafflesia* and its *Tetrastigma* hosts by identifying metabolomic shifts associated with host infection, parasitic development, and host specificity. The use of untargeted LC-MS in negative ion mode revealed distinct chemical profiles across parasitic tissues, infected and uninfected hosts, and non-host vines. These results underscore the importance of metabolite-mediated interactions in parasitic establishment and seed germination, complementing recent transcriptomic data [31] that highlight molecular mechanisms underlying host manipulation and gall-like parasitic strategies.

### 4.1 Distinct metabolomic profiles across parasitic, host, and non-host tissues

Principal component analyses revealed metabolic separation among *Rafflesia* tissues, infected and uninfected *Tetrastigma* hosts, and non-host species, indicating host metabolic reprogramming during infection. *Ampelopsis*, a putative ancestral host lineage [7], clustered separately from modern *Tetrastigma* hosts, suggesting the absence of key metabolites associated with current host compatibility. This suggests substantial chemical divergence since the ancestral host shifts inferred from phylogenomic studies.

### 4.2 Infection-associated metabolites conserved across regions

The significance scatterplot identified five metabolites consistently enriched in infected *Tetrastigma* tissues across regions, but none of these five metabolites remained significant after Benjamini–Hochberg FDR correction (*q* < 0.05); they should be interpreted as nominal, exploratory candidates rather than confirmed markers. These included one glycoside (Nb-trans-feruloylserotonin glucoside), one phytohormone/ signaling compound (Gibberellin A93), two phenolic acid/phenylpropanoids (ellagic acid and 4-methoxycinnamic acid), and one flavonoid glycoside (quercetin-3-O-deoxyhexosyl (1→2)pentoside). The predominance of phenylpropanoid- and flavonoid-derived metabolites suggests activation of defense- and stress-associated pathways, consistent with their established roles in antioxidant protection, antimicrobial defense, and signaling [32, 33, 34]. Transcriptomic data [31] similarly revealed upregulation of BURP-domain proteins, zinc-finger transcription factors, glutathione S-transferases, and genes involved in phenylpropanoid and flavonoid biosynthesis, mirroring the accumulation of phenolic metabolites in infected tissues. The enrichment of Gibberellin A93 further suggests altered gibberellin metabolism and hormonal regulation during infection. Together, these metabolites support the interpretation that infected *Tetrastigma* tissues undergo coordinated defense-related and developmental reprogramming, making them candidate chemical markers of *Rafflesia*-associated host modification.

### 4.3 Non-host *Tetrastigma* species exhibit distinct chemical defenses

Non-host *Tetrastigma* species showed distinct chemical profiles characterized by the enrichment of several phenolic metabolites. These included a metabolite putatively annotated as (Z)-resveratrol 3,4′-diglucoside, together with flavonoid- and tannin-related compounds such as epicatechin-(4β→8)-gallocatechin, (−)-epigallocatechin 3-*p*-coumaroate, and 2′,7-dihydroxy-4′- methoxy-8-prenylflavan diglucoside. Of these, only Epicatechin-(4β→8)-gallocatechin and Caffeoylmalic acid remained significant after Benjamini-Hochberg FDR correction; the remaining compounds, including (Z)-resveratrol 3,4′-diglucoside, showed nominal enrichment. Resveratrol, catechin, and broader classes of phenolics, flavonoids, and tannins have been associated with allelopathic or phytotoxic effects in other plant systems, including inhibition of seed germination and root growth [35, 36, 37]. However, such activities have not been associated with *Rafflesia* establishment and could potentially contribute to host incompatibility.

In a recent study, coumarin compounds (e.g., umbelliferone) were detected in non-host *Tetrastigma* species by LC-MS in positive ion mode (ESI+) [21]. In the present study, which employed negative ion mode (ESI-), these coumarins were not detected in non-host samples. Instead, several *p*-coumaroyl conjugates, such as (−)-epigallocatechin 3-p-coumaroate and 2-O-p-coumaroylhydroxycitric acid, were detected. These compounds and coumarins emerged within the broader phenylpropanoid metabolic network and share upstream hydroxycinnamate precursors, although the detected *p*-coumaroyl conjugates should not be interpreted as direct biosynthetic intermediates or proxies for coumarins [38]. The difference between studies may partly reflect ionization mode and other methodological differences. In a targeted analysis of eight coumarins, Ren et al. [39] reported higher-abundance precursor ions in ESI+ than in ESI- and therefore selected positive-ion mode for their analysis. Matrix-dependent ion suppression from co-eluting compounds may also reduce analyte signals under particular chromatographic and ionization conditions [40]. Consequently, the absence of coumarin signals in the present dataset should be interpreted as non-detection under the analytical conditions used rather than evidence that coumarins were absent from the tissues.

### 4.4 Metabolomic shifts across *Rafflesia speciosa* life stages and associated host tissues

The compound-level (top 20) plot shows a coherent chemical reprogramming of *Tetrastigma* during *Rafflesia* parasitism: highlighting tissue-specific enrichment such as fatty acids/oxylipins in *Rafflesia* seeds, tannins (e.g., epicatechin derivatives) in buds, elevated citric acid in infected *Tetrastigma*, and procyanidin B2 in non-host species.

*Rafflesia* seeds are enriched in fatty acids/oxylipins (e.g., 12-hydroxyoctadecanoic, palmitic, oleic), matching seed transcriptomics that indicate reliance on lipid stores via β-oxidation and gluconeogenesis and a quiescent state awaiting precise host/microbial cues [19]. Buds uniquely accumulate complex polyphenolics: (–)-catechin gallate, ellagic acid, hamamelitannin, and higher galloylated epicatechin oligomers, profiles also reported in galls and mistletoes and linked to oxidative buffering, herbivore deterrence, and potential host developmental manipulation [21, 33, 41].

Metabolomic profiling revealed that Rafflesiaceae infection across pooled *Tetrastigma* host systems is associated with reprogramming of host metabolism, characterized by enrichment of Nb-trans-feruloylserotonin glucoside, gibberellin A93, ellagic acid, 4-methoxycinnamic acid, and quercetin-3-O-deoxyhexosyl(1→2)pentoside. None of these metabolites remained significant after FDR correction and should be read as nominal, exploratory candidates. These compounds span multiple chemical classes, including phenylpropanoids, flavonoids, glycosides, and phytohormones, suggesting coordinated modulation of defense-related secondary metabolism and signaling pathways. The predominance of phenolic and flavonoid derivatives is consistent with enhanced antioxidant activity and stress-responsive metabolism commonly associated with plant biotic interactions and host-parasite relationships [42, 43, 44].

Citric acid and L-malic acid enrichment is consistent with altered central carbon metabolism and possible increased respiratory/energy demand required to support parasite sink strength, consistent with transcriptomic evidence for elevated mitochondrial activity and sugar transport [31] and with metabolic reprogramming reported in other parasitic plant systems. Host-parasite interactions are known to redirect host carbon allocation and nutrient fluxes toward the parasite, resulting in substantial metabolic changes [45, 46]. Elevated flavonoids, including eriodictyol-7- O-glucoside and epicatechin derivatives, are consistent with infection-associated phenylpropanoid remodeling and redox regulation, processes widely implicated in plant defense and stress responses [32, 48].

In contrast, non-host roots accumulated proanthocyanidins (procyanidins, cinnamtannins), phenylpropanoids (ferulic and caffeic acid derivatives, coumarin glycosides), and triterpenoids (oleanolic acid, isolimonic acid), consistent with a reinforced chemical defense phenotype [43, 44]. Procyanidin B2 enrichment may reflect a more defensive phenolic/tannin profile in non-hosts; whether this directly impedes parasite establishment remains to be tested. The accumulation of phenolics and lignification at parasite–host interfaces is a well-documented resistance mechanism against parasitic plants [48].

Most of these aerial stem/leaf-enriched compounds are phenylpropanoid-flavonoid pathway metabolites, including apigenin glycosides, isovitexin, kaempferol glycosides, phloretin derivatives, and hydroxycinnamate esters such as feruloylquinic acid. Their enrichment in aerial tissues is consistent with roles in UV protection, antioxidant defense, and stress responses rather than direct support of the belowground *Rafflesia* infection niche [49].

Collectively, these findings indicate that infected *Tetrastigma* reallocates metabolism toward energy production, signaling, and antioxidant protection, whereas non-host species show a defense-associated metabolite profile that could contribute to incompatibility, but direct functional tests are needed.

### 4.5 AI as a “Scientific Copilot” in Metabolomic Analysis

Recent advances in artificial intelligence (AI) are increasingly shaping analytical chemistry, metabolomics, and plant defense research by helping researchers interpret large and complex biological datasets. Machine learning and deep learning approaches are particularly useful for identifying biologically meaningful patterns in high-dimensional omics data [22, 23]. In plant biology, AI-assisted workflows are also becoming valuable for integrating heterogeneous datasets, identifying metabolites and biosynthetic pathways, and exploring dynamic metabolic responses involved in stress physiology and natural product biosynthesis [50]. These approaches are especially relevant in plant metabolomics, where subtle ecological and biochemical patterns often require extensive statistical processing to resolve. In the present study, AI-assisted workflows were used in a limited capacity to support Python code development, exploratory data processing, and visualization of LC-MS datasets, while all outputs and interpretations remained subject to careful expert validation and biological oversight. This approach aligns with emerging perspectives in analytical chemistry that emphasize transparency, reproducibility, and responsible human supervision in AI-supported research [25].

## 4. Conclusion

This study reveals metabolomic signatures associated with host compatibility, parasitic development, and chemical resistance in the *Rafflesia-Tetrastigma* interaction. Untargeted LC-MS shows that infection is associated with host metabolic reprogramming, bud-specific secondary metabolite production, and lipid-based readiness in seeds, while non-host Tetrastigma species showed enrichment of phenolic and flavonoid-related metabolites associated with defense or phytotoxicity in other plant systems. Together, these findings support a chemically mediated framework of parasitism in which host metabolism and signaling pathways are redirected to facilitate parasite growth and floral development. *Rafflesia* seeds may be metabolically prepared to respond to host-derived cues, although the specific signaling mechanisms remain unresolved. This work provides a molecular foundation for *Rafflesia* conservation through targeted host selection, metabolite screening, and the development of artificial infection systems that mimic permissive host environments. Generative-AI-assisted coding, together with expert oversight, supported reproducible data processing and visualization, while all scientific interpretations remained the responsibility of the authors.

## Acknowledgements

This work was made possible through the generous support of our collaborators in the Philippines. We thank our field teams, local government partners in Miag-ao and San Lorenzo Ruiz, DENR staff, and numerous colleagues and community members for their invaluable assistance. We also acknowledge contributions from Hans Banziger, Piyakaset Suksathan, and Stephen Elliott, as well as USBG, USDA, Smithsonian Gardens, and Pace University. This work is dedicated to the late Wattana Tanming and Leonard Co.

## Authors’ contributions

JM conceived the idea and led the project. JW and SP contributed to providing the permits and plant sampling materials. RA, PY, FB, and JH analyzed the data under JM’s supervision. AW, FB, PY, and MB performed the data analysis, coding validation, and visualization, and were also supervised by JM. JM, FB, and AW initiated the manuscript outlines. All authors reviewed the results and approved the final version of the manuscript.

## Data availability

The generated Python scripts are hosted on GitHub (https://github.com/adhitwicaksono/RafflesiaMetabolomics2026) along with the datasets used to generate the Figures. 1-7.

## Declarations

### Ethics approval and consent to participate

This study did not involve human participants or animals. Plant materials were obtained under National Research Council of Thailand (NRCT) sampling permit through JM and under collaboration of Queen Sirikit Botanic Garden, Chiang Mai, Thailand, and United States Botanic Garden, Washington, USA.

### Consent for publication

Not applicable.

